# A closed-loop language-model agent for target-specific multi-objective hit-to-lead optimization

**DOI:** 10.64898/2026.09.19.752921

**Authors:** Wentao Cui, Limeng Tian, Xiaoning Qi, Haoran Wang, Huanhuan Wu, Houxin He, Chen Fang, Jiaxin Hu

## Abstract

Hit-to-lead optimization is a multi-objective molecular-design problem in which binding-related scores must be considered together with drug-likeness and synthetic accessibility. We developed a closed-loop language-model medicinal-chemistry agent that combines target and starting-hit context, PubMed retrieval, molecular-property tools, docking, and feedback from previously scored analogues. In a matched computational benchmark comprising six kinase targets, six methods, a 40-candidate budget, and six independent campaigns per method, the agent achieved the highest mean aggregate score under a pre-specified composite objective on four targets. Re-analysis of the same candidate sets with drug-likeness-gated docking and docking-drug-likeness hypervolume produced different target-level winners, showing that evaluation rules can change the comparative interpretation of an optimization campaign. Candidate distributions and component ablations indicated that the agent’s advantage was associated primarily with retention of scored-candidate feedback and higher-QED regions rather than with docking alone. An EGFR case study illustrates a scaffold-preserving in-silico optimization trajectory from erlotinib. The study is a computational benchmark and evaluation framework; the prioritized molecules are hypotheses for subsequent experimental testing, not experimentally validated leads.

## 1 Introduction

Hit-to-lead (H2L) optimization converts an active starting molecule into a series of analogs whose properties support continued development. Each structural modification can alter a binding-related signal together with physicochemical properties, synthetic tractability, and medicinal-chemistry liabilities. These coupled changes make H2L a local, target-specific molecular design problem rather than a search for a single maximal score. ^1–3^ Drug-likeness and synthetic accessibility provide useful complementary views of this design space: the quantitative estimate of drug-likeness (QED) summarizes multiple properties observed in oral drugs, whereas synthetic-accessibility scores combine fragment contributions with molecular complexity. ^4,5^ A practical computational workflow therefore needs to link a defined target and starting molecule to iterative structural proposals, multi-property assessment, and a record of the outcomes of earlier proposals.

Several methodological streams provide the elements for such a workflow. Pocket-aware and retrieval-augmented models condition molecular design on protein structures and related molecular knowledge. ^6–8^ Constrained multi-property optimization and evolutionary search explore chemical spaces while accounting for competing molecular objectives. ^9–11^ In parallel, tool-augmented large language models (LLMs) can connect chemical reasoning with external information and computational operations. ^12–15^ Together, these approaches motivate an H2L agent that organizes target context, medicinal-chemistry reasoning, molecular scoring, and iterative feedback within a defined candidate budget. Its evaluation also requires an endpoint that expresses the multi-objective decision represented by the optimization task.

Here, we developed a closed-loop large language model medicinal-chemistry agent for target-aware multi-objective H2L optimization. The agent combines target and starting-hit context, pre-retrieved PubMed literature, function calls for molecular evaluation and literature search, and cumulative feedback from the highest-scoring molecules in a campaign (Fig. 1a). We paired this design with a pre-specified weak-hit protocol that selects a known inhibitor with shallow project docking for each target, thereby defining a reproducible local starting point and docking headroom (Fig. 1b). Six kinase structures, six optimization methods, six independent seeds per method, and a common 40-candidate budget establish the comparative setting. The primary aggregate score jointly represents docking, drug-likeness, synthetic accessibility, and a rule-based developability panel comprising physicochemical and structural-alert filters. We additionally report a drug-likeness-gated docking endpoint and docking-drug-likeness hypervolume, two complementary decision views of the candidate sets. ^16–18^

**Figure 1:**
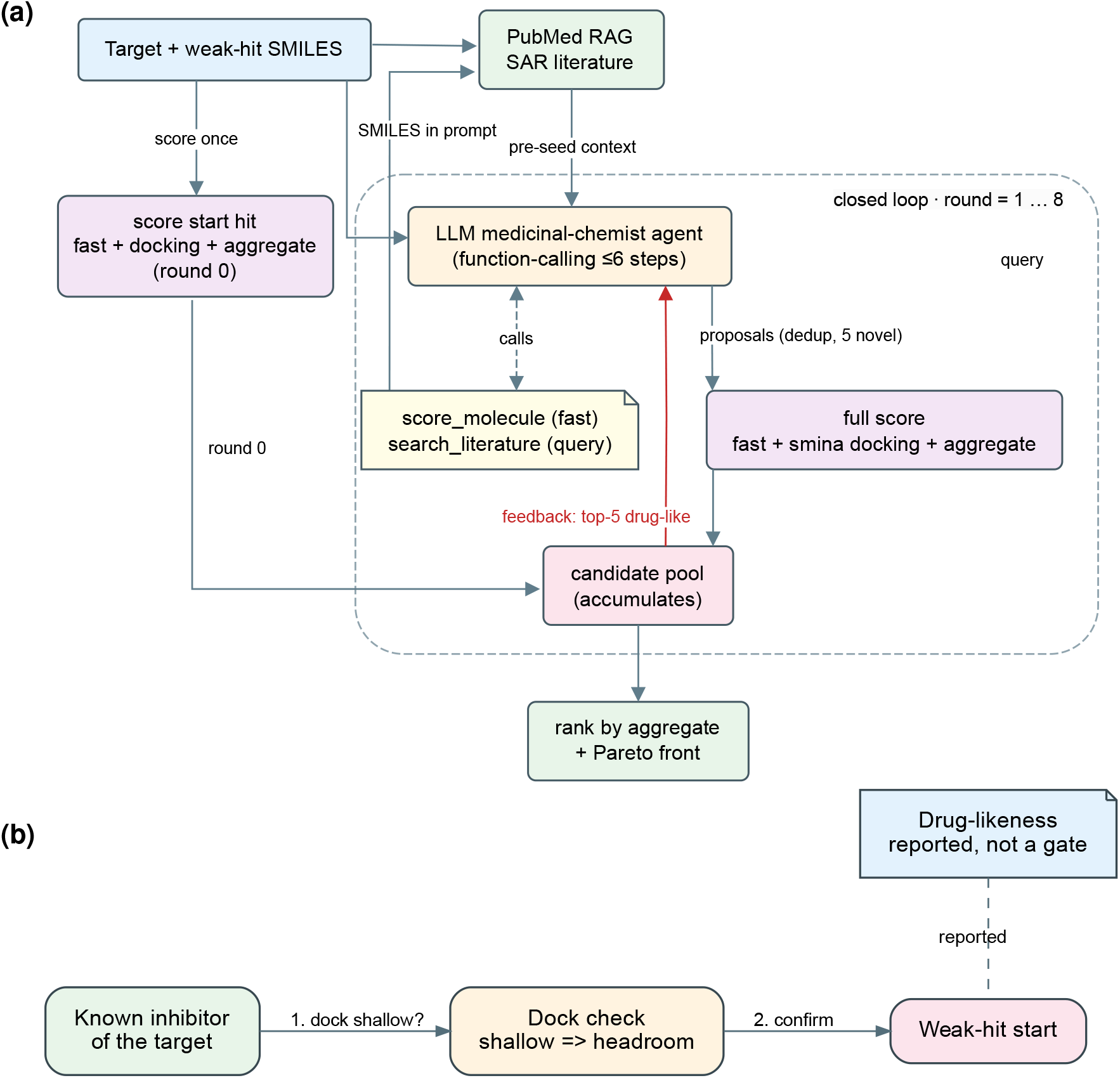
Closed-loop agent architecture and weak-hit selection protocol. (a) A target structure and starting hit are combined with pre-retrieved PubMed context in a ReAct-style medicinal-chemistry agent. The agent may query molecular-property and literature tools, proposes analogs, and receives multi-objective scores; the highest-ranked candidates provide cumulative feedback for the next round. (b) One starting compound with an established inhibitor or kinase-probe context was selected for each of six kinase structures when its project docking score left measurable optimization headroom. Quantitative estimate of drug-likeness (QED) and synthetic-accessibility (SA) score were reported properties rather than selection thresholds. The targets were EGFR (1M17), CDK2 (1AQ1), ABL1 (1IEP), BRAF (3OG7), VEGFR2 (4ASD), and SRC (3G5D).

This study provides a framework for designing and evaluating target-specific multi-objective H2L agents. We quantify aggregate performance and cumulative trajectories across six kinases, compare the winner patterns induced by the three evaluation endpoints, examine candidate docking-drug-likeness distributions and Pareto fronts, and attribute the contribution of agent components by ablation. An epidermal growth factor receptor case study further connects the closed-loop campaign to a chemically interpretable trajectory from erlotinib to prioritized analogues. By treating the headline metric as a decision rule that should match the objective coordinated during optimization, the study connects agent design, molecular-property patterns, and comparative interpretation in one target-focused H2L setting.

## 2 Results

### 2.1 A common hit-to-lead setting enabled comparison across kinase targets

We defined a matched hit-to-lead setting in which every method started from a target-specific molecule and received the same opportunity to propose new candidates. The benchmark covered six kinase structures representing distinct binding pockets: EGFR (1M17), CDK2 (1AQ1), ABL1 (1IEP), BRAF (3OG7), VEGFR2 (4ASD), and SRC (3G5D). The starting molecules were erlotinib, olomoucine, PP1, ZM336372, semaxanib, and PP2, respectively (Table 1).

**Table 1:** Target structures and pre-specified starting hits used in the six-target optimization benchmark.

| Target | PDB entry | Starting hit | Docking score<sup>a</sup> | QED<sup>b</sup> | SA score<sup>c</sup> | Developability passes<sup>d</sup> |
| --- | --- | --- | --- | --- | --- | --- |
| EGFR | 1M17 | Erlotinib | -6.80 | 0.418 | 2.48 | 2/3 |
| CDK2 | 1AQ1 | Olomoucine | -8.36 | 0.637 | 2.29 | 3/3 |
| ABL1 | 1IEP | PP1 | -7.33 | 0.744 | 2.33 | 3/3 |
| BRAF | 3OG7 | ZM336372 | -9.55 | 0.609 | 1.89 | 3/3 |
| VEGFR2 | 4ASD | Semaxanib | -7.78 | 0.737 | 2.58 | 2/3 |
| SRC | 3G5D | PP2 | -8.44 | 0.747 | 2.35 | 3/3 |

<sup>a</sup> Best smina docking affinity in kcal mol<sup>-1</sup>; lower values indicate stronger predicted binding.
<sup>b</sup> Quantitative estimate of drug-likeness.
<sup>c</sup> Synthetic-accessibility score.
<sup>d</sup> Passes across the Lipinski, Veber, and substructure-alert filters; this is a computational developability screen, not an ADMET assay.

Each starting molecule had an established inhibitor or kinase-probe context and retained measurable headroom under the study docking protocol. The starting compounds therefore defined local optimization problems with different chemical states, while the candidate budget, scoring procedure, and replicate structure were held constant. The agent connected target and starting-molecule information with literature context, property tools, multi-objective scoring, and scored-candidate feedback. This cycle generated target-specific candidate sets for subsequent comparison under a common evaluation framework.

### 2.2 The closed-loop agent led the drug-likeness-aware aggregate objective on four targets

The aggregate endpoint identified the agent as the leading method on four of the six targets: ABL1, BRAF, EGFR, and SRC (Fig. 2a). Greedy led CDK2, whereas the non-iterative LLM-blind comparator led VEGFR2. Several pairwise comparisons on the agent-leading targets were nominally significant in the replicate-level tests (Fig. 2a), whereas the remaining targets showed pocket-dependent rankings. Thus, the benchmark distinguished a recurring panel-level pattern from a universal method ordering.

**Figure 2:**
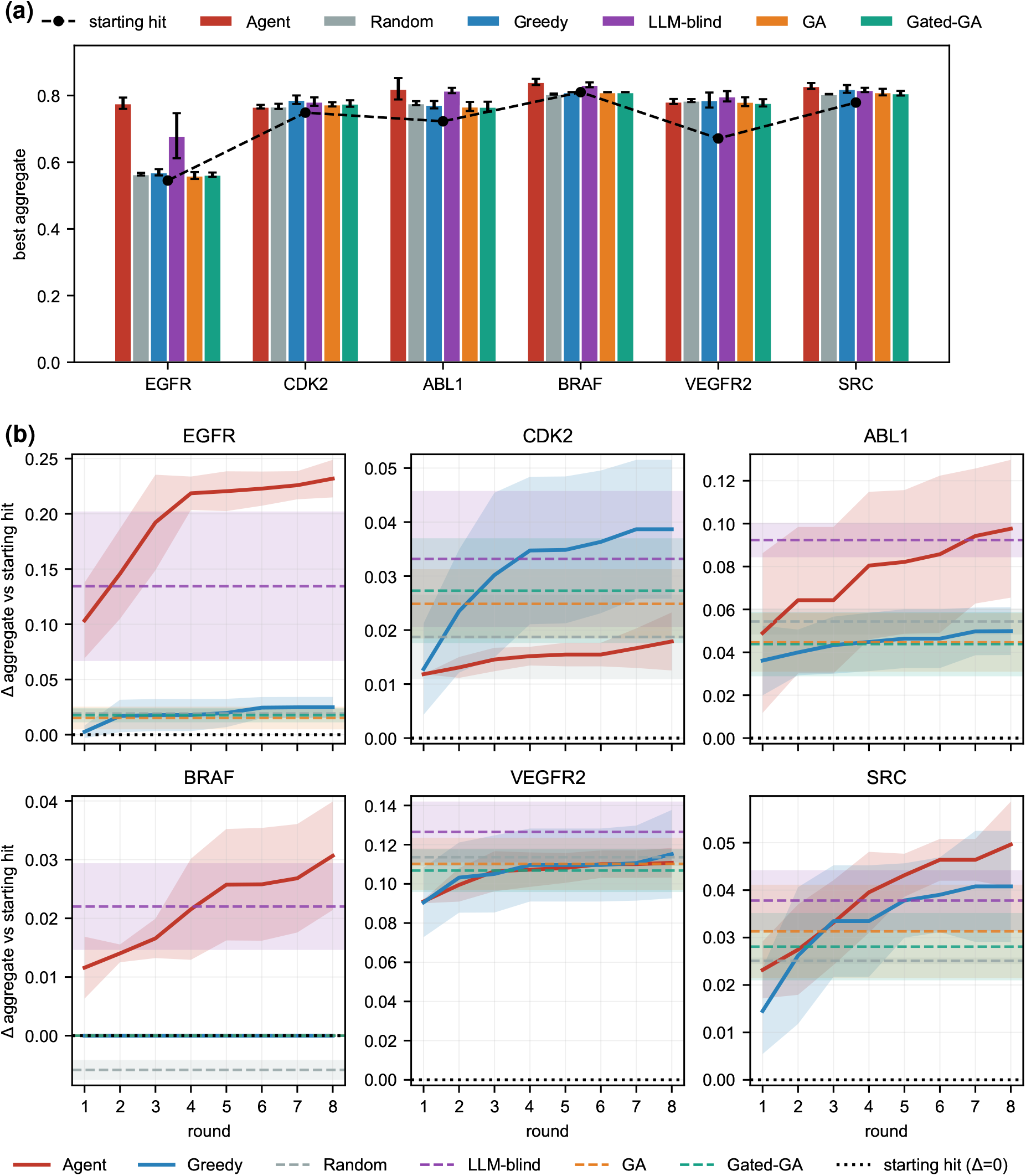
Aggregate performance and optimization trajectories across six kinase targets. (a) Mean aggregate score for six independent campaigns per method. Error bars show sample standard deviations and horizontal dashed lines show the starting-hit score. The aggregate weights docking, quantitative estimate of drug-likeness (QED), synthetic-accessibility (SA) score, and rule-based developability pass fraction by 0.35, 0.25, 0.20, and 0.20, respectively. The agent had the highest mean aggregate on ABL1, BRAF, EGFR, and SRC; greedy was highest on CDK2 and the non-iterative large language model (LLM)-blind comparator was highest on VEGFR2. On the four agent-leading targets, several one-sided Mann-Whitney U comparisons against the reference methods had nominal *P* < 0.05; values are unadjusted exploratory comparisons and each method used six campaigns. (b) Cumulative-best aggregate score by round for the agent and greedy search. Lines and shaded bands show means and sample standard deviations across six campaigns; dashed bands show the final distributions of the non-iterative comparison methods.

Round-wise cumulative-best trajectories showed how these endpoint relationships developed (Fig. 2b). On EGFR, ABL1, BRAF, and SRC, the agent progressively separated from greedy and finished with the larger aggregate value. The separation was greatest for EGFR and more modest for ABL1, BRAF, and SRC. CDK2 and VEGFR2 remained close or favored the alternative method at the final round. The trajectory pattern connected the endpoint ranking with the successive accumulation of scored molecular proposals.

### 2.3 Metric choice changed the comparative interpretation of the same campaigns

The winner depended on how the candidate set was summarized (Fig. 3a). The aggregate score assigned four target-level wins to the agent. Gated docking, which selects the best docking candidate within a QED-SA-qualified subset, distributed wins among the agent, greedy, GA, and LLM-blind methods. Docking-QED hypervolume produced a third allocation, with the agent, greedy, and LLM-blind methods each leading two targets. The same campaigns consequently supported different comparative conclusions under different decision rules.

**Figure 3:**
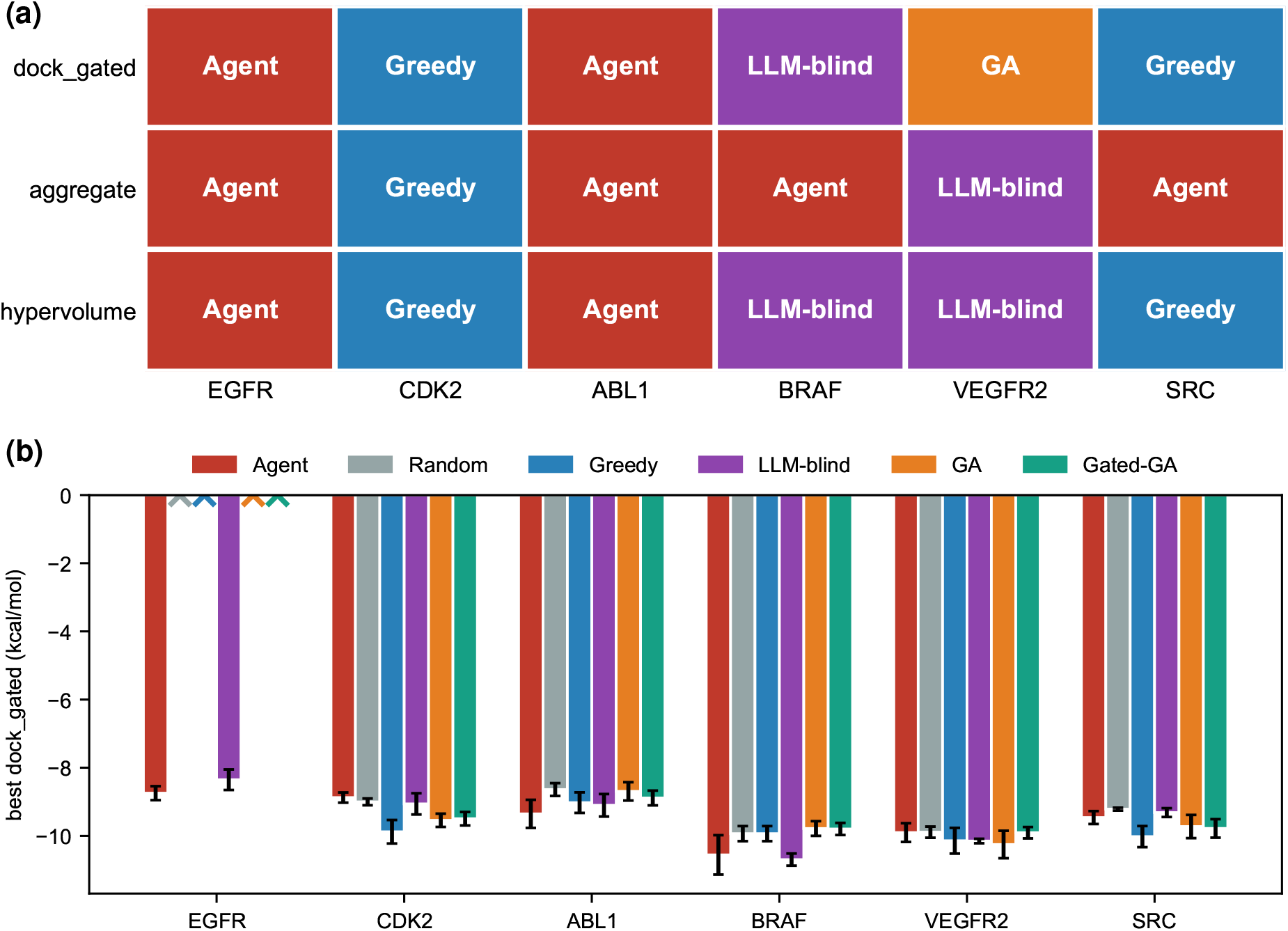
Metric alignment changes the comparative interpretation of the same optimization campaigns. The winning method for each target differs when candidate sets are summarized by aggregate score, gated docking, or docking-drug-likeness hypervolume. (b) Gated docking is the best (lowest) docking affinity among candidates meeting a quantitative estimate of drug-likeness (QED) ≥ 0.5 and synthetic-accessibility (SA) score ≤ 4; an ×denotes a method with no gate-qualified candidate across its campaigns. In the aggregate comparison, the agent was the winner for four of six targets. Under gated docking, winners were distributed among agent (two targets), greedy (two), genetic algorithm (GA; one), and large language model (LLM)-blind (one).

The target-level gated-docking panel confirmed this redistribution (Fig. 3b). The agent led EGFR and ABL1, greedy led CDK2 and SRC, LLM-blind led BRAF, and GA led VEGFR2. Because all summaries were calculated from the same candidate records, the change in winner arose from the property dimensions represented by each metric. The aggregate endpoint captured the utility optimized during H2L, whereas gated docking emphasized affinity within a restricted drug-likeness region.

### 2.4 Candidate distributions revealed the property trade-off underlying metric-dependent rankings

The combined docking-QED distributions showed that agent proposals occupied higher-QED regions while also extending towards more favorable docking scores (Fig. 4a). GA and gated-GA generated candidates in stronger-docking regions, with a larger fraction located below the QED = 0.5 reference level. These distributions provide a candidate-level explanation for the different winner patterns: movement towards stronger docking can be accompanied by a shift in QED, and the aggregate endpoint retains both dimensions in its ranking.

**Figure 4:**
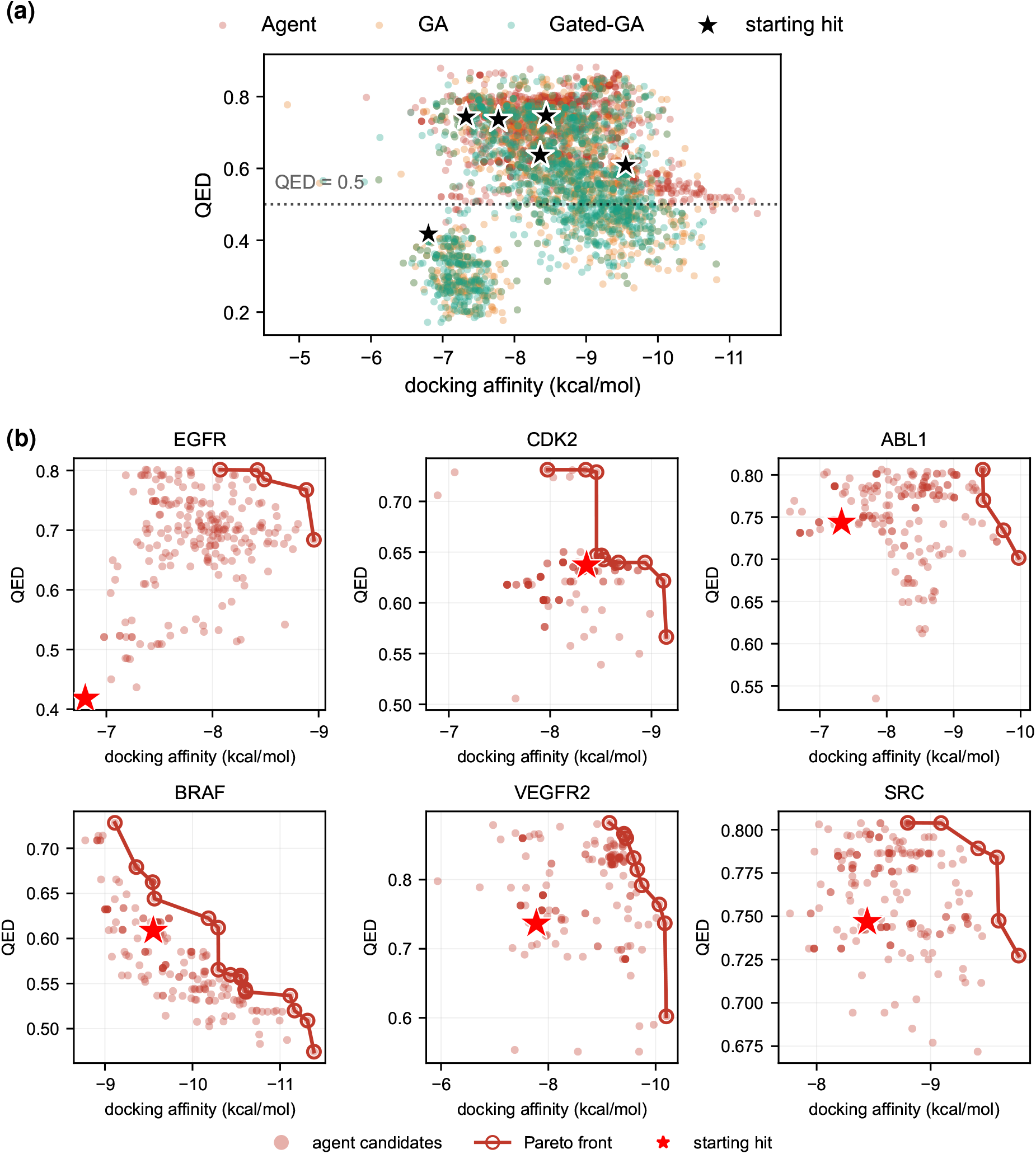
Candidate-property distributions and Pareto structure of docking affinity and quantitative estimate of drug-likeness (QED). (a) Candidate docking affinity and QED for agent, genetic-algorithm (GA), and gated-GA campaigns. The horizontal line marks QED = 0.5 and stars indicate starting hits. The agent candidate sets occupy higher-QED regions while improving docking; GA and gated-GA explore lower-QED regions associated with stronger docking. (b) Non-dominated fronts for docking affinity and QED across the six targets. These distributions provide a candidate-level explanation for the metric-dependent winner patterns in Fig. 3.

Target-resolved Pareto plots showed the same relationship in a non-dominated representation (Fig. 4b). Each agent campaign produced a front spanning the observed docking-QED trade-off from the starting hit to the prioritized candidate set. On most targets, the front extended towards stronger docking while retaining solutions in the higher-QED region; CDK2 displayed a target-specific balance along the same two-dimensional frontier. The Pareto structure linked the aggregate ranking to the distribution of candidate compromises rather than to a single molecular point.

### 2.5 Iterative scored-candidate feedback contributed most consistently to agent performance

Component ablation separated the contributions of pre-retrieved literature context, function-calling tools, and scored-candidate feedback (Fig. 5; Table 2). Removing iterative feedback lowered the best aggregate performance on five targets, with CDK2 showing the opposite direction. This broad target coverage matched the progressive separation observed in the round-wise trajectories and identified feedback as the most consistent component-level contributor.

**Table 2:** Aggregate-score effects of component removal in the agent ablation experiment.

| Target | No pre-retrieved context<sup>a</sup> |  | No tool use<sup>a</sup> |  | No iterative feedback<sup>a</sup> |  |
| --- | --- | --- | --- | --- | --- | --- |
| | $\Delta$ aggregate | *P* value<sup>b</sup> | $\Delta$ aggregate | *P* value<sup>b</sup> | $\Delta$ aggregate | *P* value<sup>b</sup> |
| EGFR | 0.0045 | 0.409 | 0.0243 | 0.197 | 0.0478 | 0.014 |
| CDK2 | -0.0120 | 0.768 | -0.0094 | 0.659 | -0.0200 | 0.999 |
| ABL1 | 0.0113 | 0.109 | 0.0076 | 0.259 | 0.0155 | 0.032 |
| BRAF | -0.0029 | 0.812 | 0.0131 | 0.001 | 0.0199 | 0.002 |
| VEGFR2 | 0.0008 | 0.531 | -0.0128 | 0.979 | 0.0101 | 0.046 |
| SRC | -0.0040 | 0.740 | 0.0102 | 0.115 | 0.0335 | 0.002 |

<sup>a</sup> $\Delta$ aggregate is the mean full-agent best aggregate score minus the corresponding ablation mean across six independent repeats; positive values indicate a higher full-agent value.
<sup>b</sup> One-sided Mann-Whitney U test comparing the full agent with the indicated ablation. Values are unadjusted exploratory comparisons.

**Figure 5:**
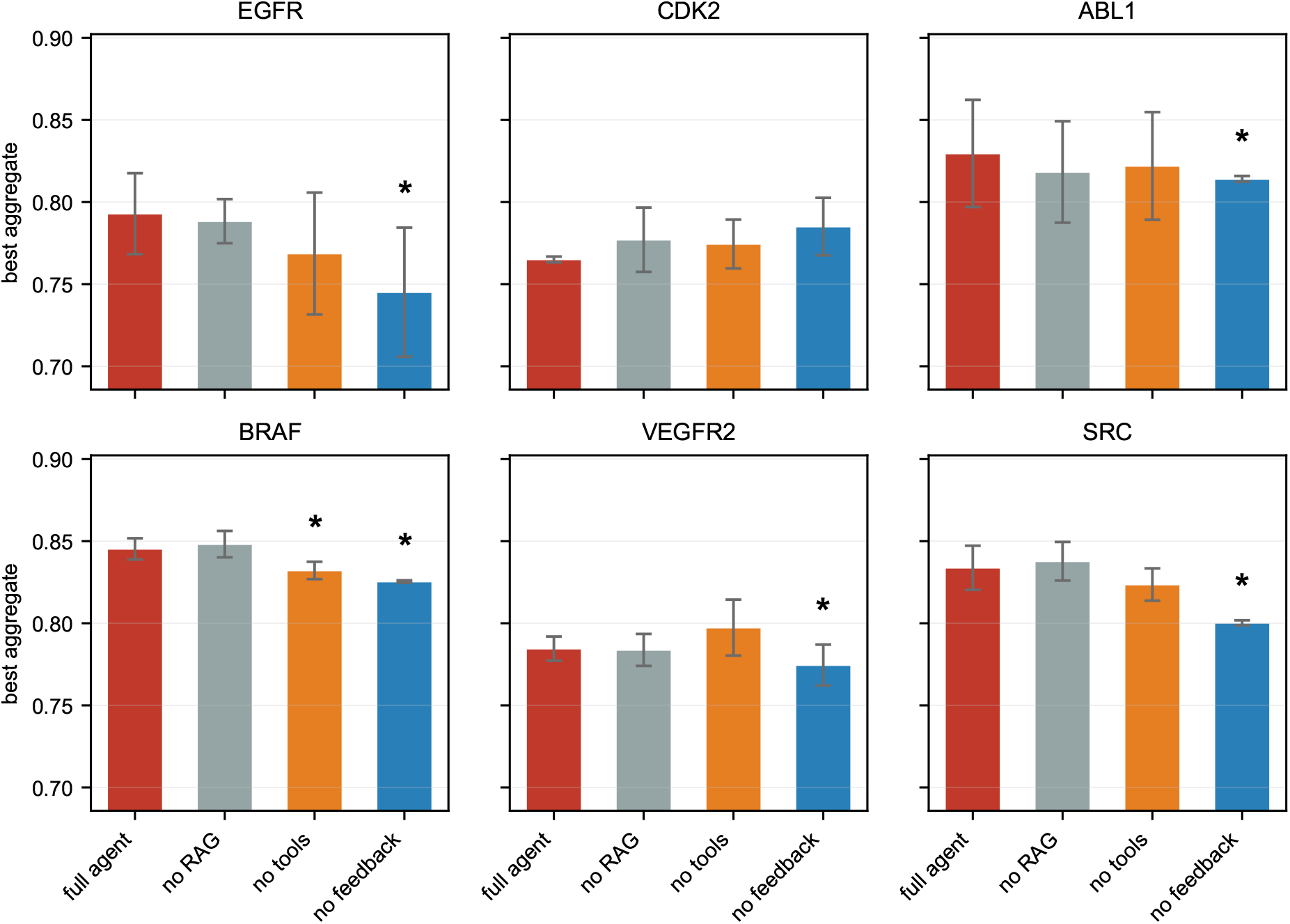
Component ablation of the closed-loop agent. The complete agent was compared with variants lacking pre-retrieved literature context (no rag), function-calling tools (no tools), or cumulative scored-candidate feedback (no feedback). Bars show the mean best aggregate score and error bars show sample standard deviations across six independent repeats. Asterisks identify one-sided Mann-Whitney U comparisons with nominal *P* < 0.05 against the full agent. Removing iterative feedback reduced aggregate performance on EGFR, ABL1, BRAF, VEGFR2, and SRC; removal of pre-retrieved literature context produced no nominal target-level comparison.

The other ablations produced more local changes. Aggregate performance remained comparable across targets when the pre-retrieved context was removed, while tool removal produced a clear target-level change only for BRAF. The no rag comparison retained on-demand literature search and therefore isolated the context supplied before generation. Together, the ablation results localised the principal contribution of the closed loop to the reuse of scored candidates between rounds.

### 2.6 An EGFR campaign illustrated coordinated molecular changes

We followed one EGFR campaign from erlotinib to its final agent analogue to examine the molecular form of the aggregate improvement (Fig. 6). A representative seed-0 trajectory passed through intermediate structures before reaching the round-8 endpoint. The 4-anilinoquinazoline core was retained, while peripheral substituents were modified, yielding a scaffold-preserving analog path.

**Figure 6:**
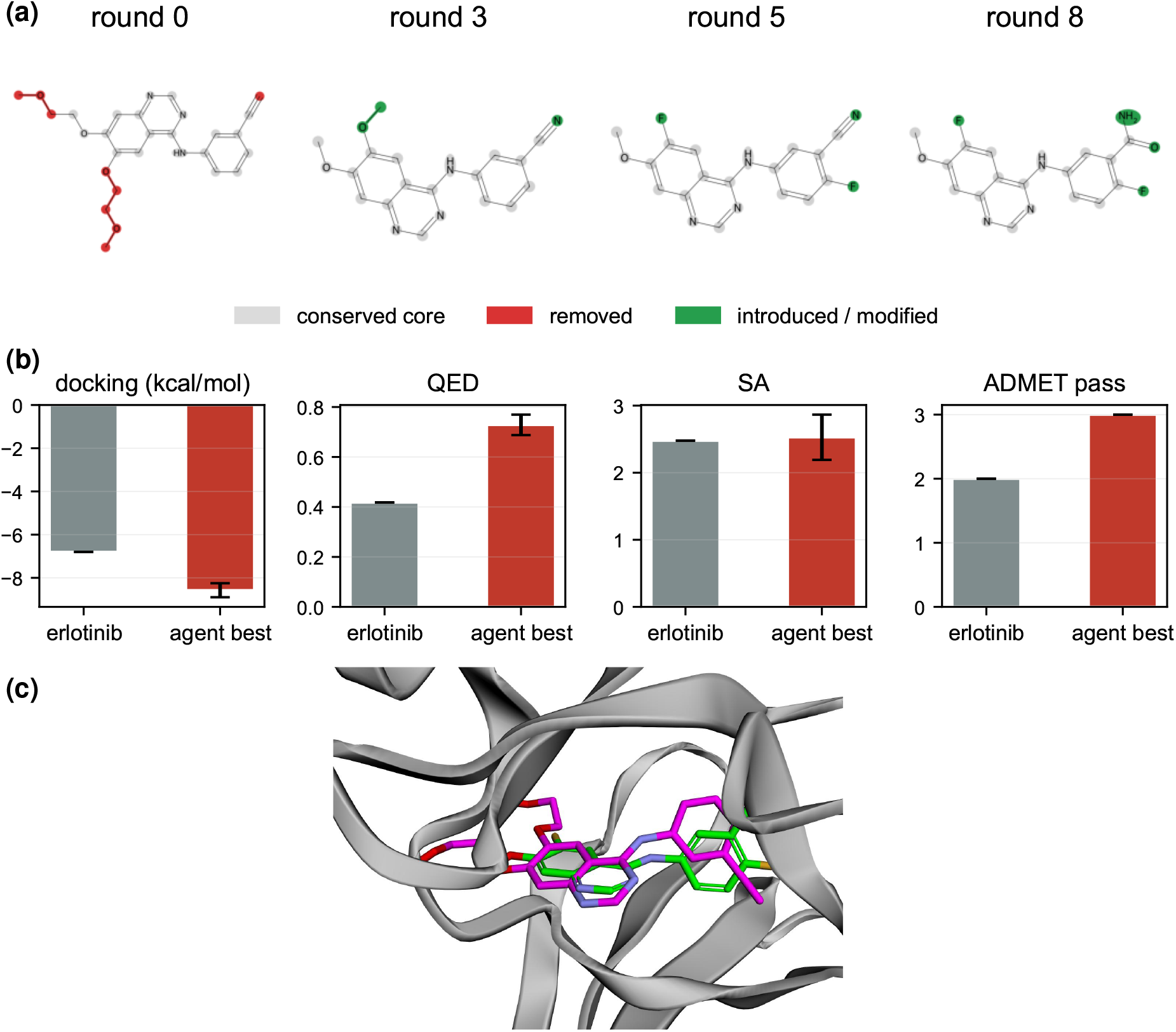
EGFR molecular-design trajectory from erlotinib to a prioritized analogue. (a) A representative seed-0 trajectory through rounds 0, 3, 5, and 8. The 20-atom 4-anilinoquinazoline maximum common substructure is shown in gray; removed atoms are red and introduced or modified atoms are green. (b) Erlotinib and the best agent analog across six seeds. Docking affinity changed from -6.80 to −8.57 ± 0.32 kcal mol^-1^, quantitative estimate of drug-likeness (QED) from 0.418 to 0.729 ± 0.041, synthetic-accessibility (SA) score from 2.48 to 2.53 ± 0.34, and rule-based developability passes from 2/3 to 3/3. (c) Superposition of the docked endpoint analog (green) and co-crystal erlotinib (magenta) in the ATP-binding pocket of EGFR structure 1M17.

Across six seeds, the best agent analogue improved the docking score from the erlotinib starting value of -6.80 kcal mol^-1^ to approximately -8.6 kcal mol^-1^. QED increased from 0.418 to approximately 0.73, the SA score remained at a similar level, and the developability-panel pass count increased from 2/3 to 3/3 (Fig. 6b). The case therefore recapitulated the benchmark-level pattern of stronger docking accompanied by improved or retained drug-likeness-related properties. The docked endpoint analogue superposed with the co-crystal erlotinib pose in the ATP-binding pocket of EGFR 1M17 (Fig. 6c), providing a structural plausibility check for the retained core and peripheral modifications.

### 2.7 Orthogonal checks contextualised the docking readout

Three-seed rescoring of the pooled benchmark molecules showed generally stable docking scores and high agreement in molecular ranking, with a narrower score range and lower rank stability for VEGFR2. Co-crystal redocking reproduced the deposited pose within 2 °A for four of the six targets; BRAF and SRC showed larger deviations. These assessments defined the target-specific context for interpreting docking-based comparisons and the EGFR pose (Table 3).

**Table 3:**
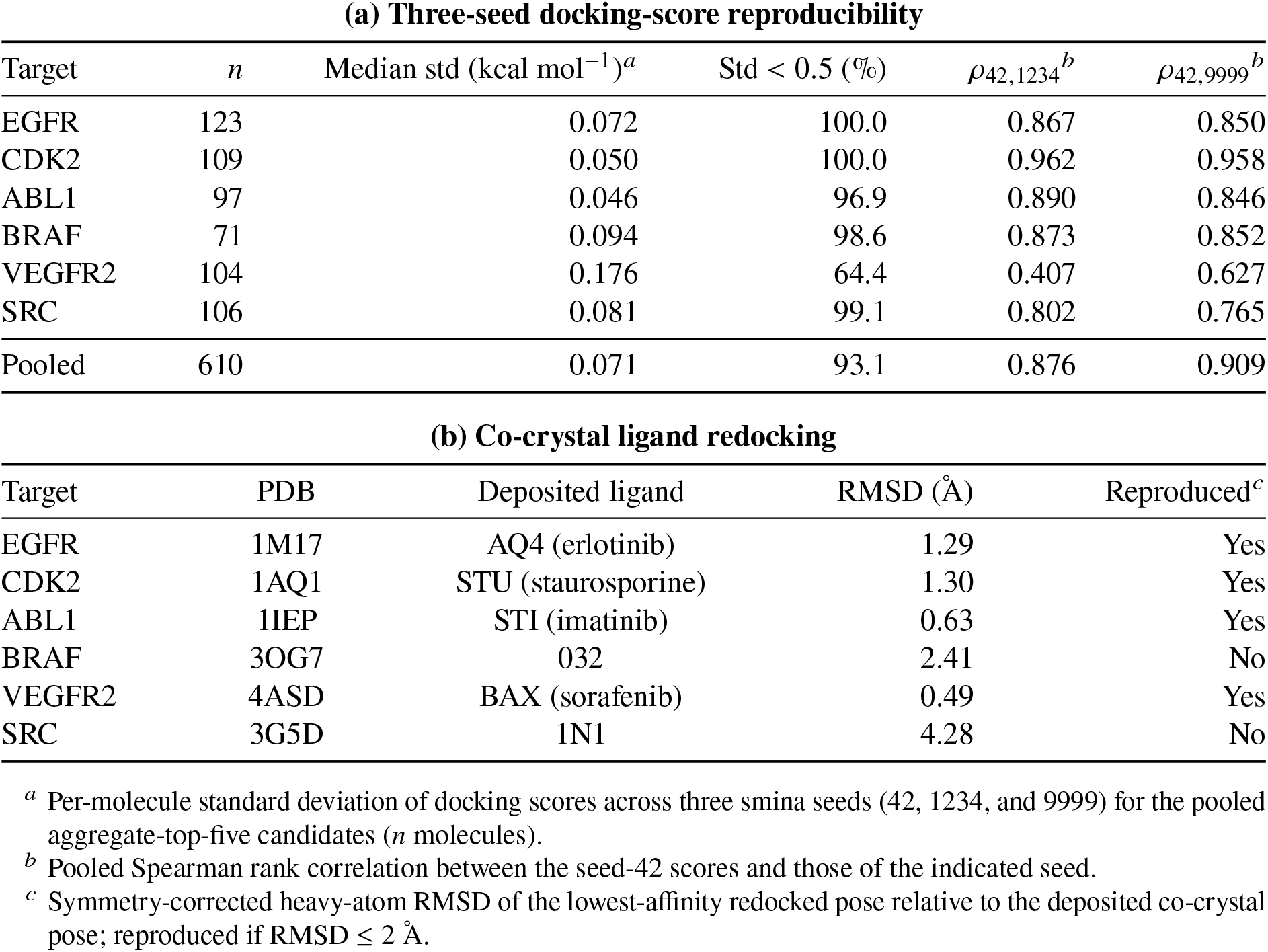
Orthogonal validation of the docking readout.

## 3 Discussion

Hit-to-lead optimization requires decisions over a set of molecular properties rather than over a single predicted attribute. ^1,2^ In the six-kinase, equal-budget benchmark, the closed-loop agent achieved the highest mean aggregate score on four targets, while the method winning under docking among drug-like candidates and under docking-drug-likeness hypervolume varied by target. The docking-QED distributions and the EGFR trajectory placed these aggregate results at the molecule level, and the ablation experiment identified scored-candidate feedback as the component with the most consistent contribution. Taken together, these observations show how a target-conditioned agent can coordinate the computational objective defined for a local hit-to-lead campaign, and they identify metric alignment as important to interpreting its output.

The trajectory and ablation results suggest that the agent’s operational contribution arose from coupling structural proposals to the accumulated record of multi-property outcomes. ReAct provides a general pattern for interleaving reasoning and actions, whereas chemistry-oriented systems have shown how language models can be grounded in tools and chemical information. ^12–14^ Structure-aware molecular generation and optimization methods likewise condition candidate design on protein or molecular context. ^6–8,15^ The present workflow anchors those capabilities to a defined target and starting hit, then reuses the highest-ranked scored analogs at each round under a shared candidate budget. The higher cumulative aggregate trajectories on four targets and the high-QED candidate regions are consistent with this feedback retaining information about both binding-related and developability-oriented objectives. Removing pre-retrieved literature context did not change aggregate performance at the target level in this experiment; because on-demand literature search remained available, this result specifically describes the contribution of the pre-retrieved context channel in the implemented workflow.

The comparison also illustrates a general principle for multi-objective molecular design. An optimization method produces a candidate set, and each reported metric applies a different decision rule to that set. Here, the aggregate score represents the four-component utility used during optimization, *dock gated* represents docking within a QED-SA-qualified subset, and hypervolume represents the extent of the dockingdrug-likeness Pareto region. ^3,17,18^ Their distinct winner patterns therefore describe complementary decisions rather than interchangeable estimates of a single performance quantity. The aggregate advantage, distributed *dock gated* winners, and docking-QED patterns together support reporting a primary endpoint that represents the design objective, alongside complementary decision views and the candidate-property distribution that generated them. This approach extends established molecular-design benchmarking and Pareto analysis by making the relationship between the optimization objective and the headline conclusion explicit. ^10,11,16^

The current evidence covers six kinase structures, one pre-specified starting hit per target, six independent campaigns per method, a 40-candidate budget, and four computational scoring components. Three-seed rescoring of 610 molecules produced a median per-molecule docking-score standard deviation of 0.071 kcal mol^-1^, with 93.1% below 0.5 kcal mol^-1^ and pooled rank correlations of 0.876 and 0.909; VEGFR2 showed a narrower score range and lower rank stability. Co-crystal redocking reproduced four of six structures within 2 °A (Table 3), with larger deviations for the BRAF and SRC ligands (2.41 and 4.28 °A). These assessments give a target-specific context for the docking component, whose interpretation in virtual screening depends on pose and scoring behavior. ^19,20^ The study does not establish biochemical activity, cellular efficacy, selectivity, synthetic feasibility or experimental ADME. The prioritized molecules should therefore be interpreted as computational hypotheses, and the generality of the framework beyond this six-kinase benchmark remains to be tested.

Several computational limitations define the scope of the conclusions. The composite objective and the QED-SA gate were specified by the study and are not universal measures of lead quality; because the agent was prompted with related objectives, part of its advantage is necessarily objective-specific. The six targets, one starting hit per target and six repeats per method provide a controlled benchmark but do not support broad claims across chemical space. Docking-score reproducibility was also target-dependent, and co-crystal redocking deviated substantially for BRAF and SRC. Future computational work should therefore test sensitivity to objective weights and gates, include independent scoring functions, use proposal-space-matched baselines and evaluate additional targets and starting points.

## 4 Methods

### 4.1 Target structures and weak-hit starting points

We established target-specific hit-to-lead campaigns for six kinase structures deposited in the Protein Data Bank (PDB) ^21^: epidermal growth factor receptor (EGFR, 1M17), cyclin-dependent kinase 2 (CDK2, 1AQ1), Abelson tyrosine kinase 1 (ABL1, 1IEP), B-Raf proto-oncogene (BRAF, 3OG7), vascular endothelial growth factor receptor 2 (VEGFR2, 4ASD), and SRC kinase (SRC, 3G5D). For each structure, the receptor file, cognate-ligand chain and residue identifier, and docking-box center and dimensions were recorded in a target-specific metadata file. The six starting molecules were erlotinib for EGFR, olomoucine for CDK2, PP1 for ABL1, ZM336372 for BRAF, semaxanib for VEGFR2, and PP2 for SRC. These molecules were selected before optimization by combining a known inhibitor or kinase-probe context with a relatively shallow score from the study docking protocol (Fig. 1b), thereby defining a tractable local optimization scenario. Their starting docking scores, QED values, and synthetic-accessibility (SA) scores are reported in Table 1. The inhibitor contexts were drawn from the original studies of erlotinib, semaxanib, ZM336372, olomoucine, and PP2. ^22–26^ QED and SA were recorded as starting-point properties rather than used to exclude a starting molecule.

### 4.2 Closed-loop large language model agent

The closed-loop agent received the target name, its co-crystal ligand, and the starting-hit Simplified Molecular Input Line Entry System (SMILES) string. Its system instruction specified the role of a medicinal chemist and requested chemically valid analogs that jointly improve binding-related docking score, QED, SA, and a developability panel. The implementation followed a reasoning and acting (ReAct) pattern in which the model could interleave molecular proposal with tool calls ^12,13^ (Fig. 1a). Before a campaign, the agent queried PubMed for “*target* inhibitor structure-activity relationship”, retrieved up to eight records through the NCBI E-utilities interface ^27^, and inserted their title-abstract snippets into the model context. Retrieved records were ranked with BM25. ^28^ During generation, the model could `call_score molecule` for rapid RDKit property and developability-panel calculations and `search_literature` for an on-demand PubMed query; a maximum of six tool-interaction steps was permitted per proposal call. The reported campaigns used the model identifier `gpt-5.6-luna` through an OpenAI-compatible endpoint at temperature 0.7; prompts, tool schemas, configuration and raw candidate records are provided in the public repository.

The initial hit was scored at round 0. Each subsequent round requested five analogs, removed invalid or previously observed SMILES strings, and evaluated the remaining proposals. The prompt for the next round contained up to five previously scored molecules that satisfied QED ≥ 0.5 and SA ≤ 4, ordered by docking score, together with their four-property feedback. Campaigns comprised eight rounds and five requested proposals per round, corresponding to a 40-proposal optimization schedule. Candidate records retained the proposed SMILES string, round, origin, docking score, molecular properties, and aggregate score.

### 4.3 Molecular scoring and multi-objective summaries

All candidates were first processed with RDKit. ^29^ We calculated QED ^4^, the fragment- and complexity-based SA score ^5^, molecular weight, calculated logP, topological polar surface area, hydrogen-bond donors and acceptors, rotatable bonds, and ring counts. Candidates with an invalid SMILES string or non-finite QED or SA value were excluded before docking. The developability panel comprised three binary checks: Lipinski compliance (molecular weight ≤500, logP ≤5, hydrogen-bond donors ≤5, and hydrogen-bond acceptors ≤10) ^30^; Veber compliance (rotatable bonds ≤10 and topological polar surface area ≤140) ^31^; and absence of RDKit PAINS, Brenk and NIH structural alerts. ^32,33^ The resulting panel score was the number of passed checks divided by three. This rule panel is a computational developability screen and is not an experimental or predictive ADMET assay.

Docking used smina, an AutoDock Vina derivative ^34,35^, with the target-specific receptor and box coordinates. A three-dimensional ligand was embedded from SMILES using RDKit with seed 42, optimized by the Merck molecular force field (MMFF) when that calculation succeeded, and written as a structure-data file (SDF). Smina was run at exhaustiveness 8 with four CPU threads; the lowest affinity parsed from the output poses was retained in kcal mol^-1^. Parallel candidate evaluation used up to 32 worker processes. A lower docking affinity was treated as favorable.

The primary aggregate score was calculated as

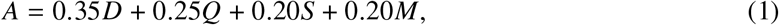

where *D* is docking affinity linearly mapped from -4 to -11 kcal mol^-1^ and clipped to [0, 1], *Q* is QED, *S* = (10 −SA /9) clipped to [0, 1], and *M* is the developability-panel pass fraction. Higher *A* values are favorable. The weights and thresholds were pre-specified for this benchmark and are not universal measures of lead quality. We also calculated three complementary candidate-set summaries. *Dock gated* was the most favorable docking score among candidates with QED ≥0.5 and SA ≤4. The non-dominated set was determined over raw docking affinity (minimized) and a normalized non-docking drug-likeness blend comprising QED, SA, and the developability-panel score. Its two-dimensional hypervolume used the reference point (−4, 0) after transforming docking so that both axes increase with improvement. ^17^ Chemical diversity was the mean pairwise 1 Tanimoto distance from radius-2, 2,048-bit Morgan fingerprints. ^36^

### 4.4 Comparator methods

Five comparator methods used the same candidate-property evaluator and target metadata. The random baseline generated independent single-site analogs of the starting hit by attaching one fragment selected from F, Cl, Br, cyano, trifluoromethyl, hydroxy, amino, methoxy, methyl, or cyclopropyl substituents. The greedy baseline generated the same class of analogs over eight rounds and selected the highest aggregate-scoring molecule as the parent for the following round. The *llm blind* baseline used the same language-model service to generate a single batch of analogs from the target and starting hit, without iterative property feedback or tool calls.

The genetic-algorithm (GA) baseline implemented a (*µ* +*λ*) evolution strategy with a population of eight molecules, mutation by the same single-site operator, and elitist selection by aggregate score. ^37^ It ran until the 40-candidate budget was reached. The gated-GA used the same population procedure, retained candidates satisfying QED ≥ 0.5 and SA ≤4 for parent selection, and selected these parents by docking affinity. These mutation- and evolutionary-search comparators provide established molecular-optimization reference strategies. ^10,11,38^

### 4.5 Benchmark, component ablation, and statistical analysis

The benchmark crossed six targets, six methods (agent, random, greedy, *llm blind*, GA, and gated-GA), and six completed campaign replicates indexed 0-5. The agent and *llm blind* replicates reflected independent stochastic language-model calls; the random, greedy, and GA procedures used the corresponding pseudorandom-number-generator index. The agent and greedy procedures used the eight-round, five-proposal schedule, while the other comparators used a nominal 40-candidate allocation. Candidate sets were summarized by aggregate score, *dock gated*, hypervolume, and diversity. For each target-metric comparison, the agent was compared with each baseline by a one-sided Mann-Whitney *U* test on the six replicate-level values. ^39^ The alternatives were agent > baseline for aggregate score, hypervolume, and diversity, and agent < baseline for *dock gated*. Means are reported with sample standard deviations; *P* < 0.05 denotes an unadjusted exploratory one-sided comparison.

Component ablations used the same six targets, eight rounds, five-proposal schedule, scorer, and six independent repeats. The full configuration combined pre-retrieved PubMed context, function calling, and scored-molecule feedback. The *no rag* variant used target and starting-hit context with on-demand `search literature` access, supplying literature during generation through that tool. The *no tools* variant generated proposals directly from the target and starting-hit context, after which all candidates underwent evaluation with the common scorer. The *no feedback* variant retained literature and tool access, while subsequent prompts drew on the target and starting-hit context rather than scored candidates. Each variant was compared with the full agent using a one-sided Mann-Whitney *U* test, with the alternative hypothesis that the full agent had a higher best aggregate score per repeat.

### 4.6 Docking-score consistency and pose assessment

We performed two orthogonal assessments of the docking component, whose interpretation requires attention to both scoring and pose accuracy. ^19,20^ First, the aggregate-top-five candidates from every target, replicate, and method were pooled and deduplicated, yielding 610 molecules. Each was redocked with smina seeds 42, 1234, and 9999 at the same target-specific box, exhaustiveness, and CPU settings. We calculated the per-molecule standard deviation across the three scores and pooled Spearman rank correlations between seed 42 and each of the other seeds.

Second, each deposited co-crystal ligand was redocked into the box used for its target campaign. The lowest-affinity smina pose was compared with the deposited ligand pose after removing hydrogens, using the symmetry-corrected heavy-atom root-mean-square deviation (RMSD) implemented in RDKit.

## Data availability

The protein structures used in this study are publicly available from the Protein Data Bank under accessions 1M17, 1AQ1, 1IEP, 3OG7, 4ASD and 3G5D.

## Code availability

MedChemAgent source code, analysis scripts, computational-environment specification and implementation configuration are available at https://github.com/Hjxzuibang/MedChemAgent under an MIT license.

## Author contributions

W.C. contributed to funding acquisition, resources, investigation, validation, data curation and writing-review and editing. L.T. contributed to software, validation and visualization. X.Q. contributed to methodology, validation, investigation, resources and writing-review and editing. H.Wa. contributed to software, validation and data curation. H.Wu. contributed to validation, investigation and data curation. H.He. contributed to validation, resources and writing-review and editing. C.F. contributed to supervision, project administration, resources and writing-review and editing. J.H. contributed to conceptualization, methodology, software, formal analysis, investigation, data curation, validation, visualization, writing-original draft, writing-review and editing, supervision and project administration. All authors approved the final manuscript.

## Acknowledgements

This work was supported by Beijing Life Science Academy (2025900CC0280, 2025900CB0170). The language model gpt-5.6-luna was used as the computational proposal-generation component; it is not an author, and all study design, analyses and manuscript decisions were made by the authors.

